# A Protein Language Model for Exploring Viral Fitness Landscapes

**DOI:** 10.1101/2024.03.15.584819

**Authors:** Jumpei Ito, Adam Strange, Wei Liu, Gustav Joas, Spyros Lytras, The Genotype to Phenotype Japan (G2P-Japan) Consortium, Kei Sato

## Abstract

Successively emerging SARS-CoV-2 variants lead to repeated epidemic surges through escalated spreading potential (i.e., fitness). Modeling genotype–fitness relationship enables us to pinpoint the mutations boosting viral fitness and flag high-risk variants immediately after their detection. Here, we introduce CoVFit, a protein language model able to predict the fitness of variants based solely on their spike protein sequences. CoVFit was trained with genotype–fitness data derived from viral genome surveillance and functional mutation data related to immune evasion. When limited to only data available before the emergence of XBB, CoVFit successfully predicted the higher fitness of the XBB lineage. Fully-trained CoVFit identified 549 fitness elevation events throughout SARS-CoV-2 evolution until late 2023. Furthermore, a CoVFit-based simulation was able to predict the higher fitness of JN.1 subvariants before their detection. Our study provides both insight into the SARS-CoV-2 fitness landscape and a novel tool potentially transforming viral genome surveillance.

## Introduction

A primary challenge faced in controlling viral infectious diseases stems from the ability of viruses to evolve through mutations^1^. Throughout the COVID-19 pandemic, SARS-CoV-2 variants with escalated spreading potential (i.e., fitness) in the host population have successively emerged, leading to repeated epidemic surges^2,3^. By understanding how viruses enhance their fitness in a pandemic through the lens of SARS-CoV-2 studies, we can learn critical insights for managing not just COVID-19 but future viral infectious diseases as well.

The fitness of SARS-CoV-2 variants can be compared using the effective reproduction number (R_e_)^4–7^. R_e_ represents the average number of secondary infections caused by an infected individual in a certain condition. Owing to advancements on virus genome surveillance, it is now feasible to estimate the relative R_e_ of SARS-CoV-2 variants in almost real-time. Based on the estimated relative R_e_, we can predict which variant(s) has the biggest advantage amongst co-circulating viruses at a given time and will likely become the next dominant variant. Our team, the G2P-Japan consortium, has successfully used R_e_ estimation to predict upcoming dominant variants and further elucidated the characteristics and risks these variants possess through virological experiments^8–20^.

Since the fitness of a variant in a certain condition is determined by its genotype, it is feasible, in principle, to predict the fitness of a variant from its genome or protein sequences. By establishing a fitness prediction model, we can identify mutations that contribute to increased viral fitness. Furthermore, a reliable fitness prediction model would enable an efficient variant monitoring system, ideally capable of identifying the next dominant variants as soon as the initial genome sequence of them is made available. Moreover, since viruses typically evolve in a direction that increases their fitness, understanding the virus’s fitness landscape enable to predict their evolution.

SARS-CoV-2 enhances its fitness by acquiring mutations (including substitutions, insertions, and deletions) in viral proteins, with particular emphasis on the spike (S) protein^6^. The S protein is the glycoprotein essential for virus entry into the host cells via interaction with the angiotensin-converting enzyme 2 (ACE2) receptor^21^. Also, the S protein is a primary target for neutralizing antibodies (Abs), which are key components of the humoral immune response triggered by vaccinations or natural infections^22^. Therefore, mutations in the S protein that can affect its binding efficiency with ACE2 and its ability to evade neutralizing Abs tend to have stronger impact on viral fitness^2,3,8–20^.

The evolutionary process of SARS-CoV-2 can be categorized into sequential and non-sequential (or saltation-like) evolution^2,3^. Omicron BA.1, BA.2, XBB, and BA.2.86 lineages emerged through a saltation-like process, acquiring greater than approximately 15 mutations in the S protein through various mechanisms including recombination^8,9,13,19,23^. In contrast, Omicron BA.5, BQ.1.1, and descendants of XBB evolved sequentially from their parental lineages, acquiring a smaller number of mutations in the S protein, typically three or fewer^10,12,14–16,18^. Predicting the fitness of variants emerged through the saltation-like evolution is expected to be more challenging than the fitness of variants emerged through sequential evolution.

Previous studies, including ours, have developed methods to predict the fitness of variants based on their mutation patterns using a statistical modeling approach^6,12,24,25^. However, these statistical models simply represent fitness as a linear combination of individual mutation effects, not considering interactions between mutations, namely epistasis^6,12,24,25^. Furthermore, these statistical models could not consider the effect of mutations that have not emerged as of the training dataset creation. We assumed that these challenges can be addressed using innovative protein language models^26^. Large language models (LLMs) have become increasingly popular in the field of natural language processing. Protein language models are LLMs that have been pretrained on comprehensive datasets of protein sequences, enabling them to learn the general patterns of proteins. Notably, we can finetune a pretrained protein language model to solve specific tasks, like fitness prediction. Moreover, by employing a multitask learning framework, applicable for deep learning models such as protein language models, we can integrate functional information on mutations into the fitness prediction process, potentially enhancing the model’s predictive accuracy.

In this study, we developed CoVFit, a model to predict the fitness of SARS-CoV-2 variants based on the S protein by utilizing the state-of-the-art protein language model ESM-2^27^. We finetuned a customized ESM-2 model using i) genotype–fitness information, estimated from virus genome surveillance, and ii) mutation effect information on evasion ability from humoral immunity, determined by high-throughput deep mutational scanning (DMS) experiments^28^. Furthermore, using CoVFit, we have depicted the fitness landscape of SARS-CoV-2 until late 2023.

## Results

### Introduction of CoVFit

We developed CoVFit, a fitness prediction model based on S protein sequences by finetuning ESM-2 (**Fig. 1A**). To increase the model’s knowledge of the coronavirus S proteins, we first established ESM-2_Coronaviridae_, by performing additional pre-training (i.e., domain adaptation) on the ESM-2 model with S protein sequences obtained from 1,506 *Coronaviridae* viruses (**Fig. 1B** and **Fig. S1**). The ESM-2_Coronaviridae_ model demonstrates enhanced predictive capability in the masked learning task specifically for SARS-CoV-2 S proteins, while retaining its performance across a broader collection of proteins (**Fig. S1C**). Subsequently, utilizing a multitask learning framework, we finetuned the model on both genotype–fitness (R_e_) data and DMS data for the ability to escape neutralization by Abs (**Fig. 1B**). Consequently, for a given S protein sequence, CoVFit can predict the country-specific fitness value and the ability to escape from each mAb (**Fig. S1A**).

**Fig. 1.**
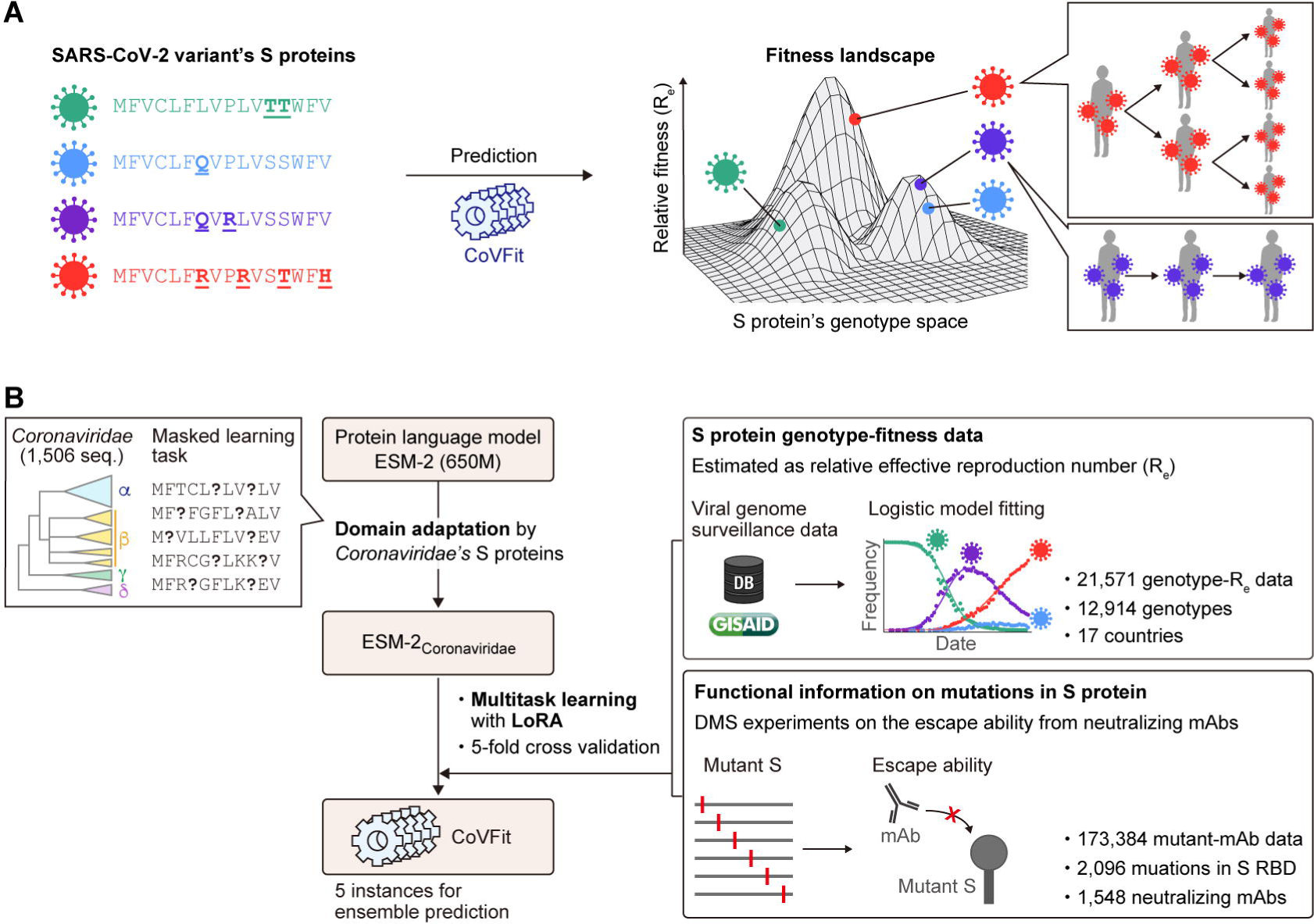
Overview of CoVFit. A) Conceptual framework of CoVFit. CoVFit is a protein language model designed to predict the relative fitness (R_e_) of SARS-CoV-2 variants based on their S protein sequences. B) Outline of the training process used to develop CoVFit model instances.

To assemble the genotype–fitness dataset, we first classified viral sequences into S protein genotypes, defined as groups of viruses sharing a unique set of mutations in S protein. Subsequently, we estimated the R_e_ of each genotype in each country by fitting a multinomial logistic model to the genome surveillance data up to November 2, 2023, obtained from GISAID (https://gisaid.org/), as previously described^2,3,8–20^. Consequently, we obtained a total of 21,751 genotype–fitness data points, covering 12,914 genotypes across 17 countries (**Figs. 1B and S2A**). Consistent with a previous study^6^, there is a clear trend where later-emerging variants exhibit higher R_e_ values, suggesting a gradual increase in the R_e_ of variants through evolution (**Fig. S2A**).

We utilized an *in vitro* DMS dataset on the neutralization capabilities of monoclonal Abs (mAbs), produced by Cao et al.^29^. A total of 173,384 mutation– mAb DMS data points, covering 2,096 types of mutations in the receptor binding domain (RBD) in the S protein and 1,548 mAbs, were included in the dataset (**Figs. 1B and S2B**). Aligning with previous findings^29^, the effects of mutations on mAbs varied depending on their epitope classes **Figs. S2B**, **S2C, and S2D**). Variants exhibiting higher fitness, such as BQ.1.1 or XBB, exhibited an increased ability to evade these mAbs, supporting the effectiveness of utilizing this information for predicting fitness (**Fig. S2D**).

Using the five-fold cross validation scheme, we generated five model instances of CoVFit (**Fig. 1B**). These model instances were used to evaluate performance for the corresponding test data. Additionally, these instances can provide the mean and variance of predictions for new data (e.g., data acquired after model training). Hereafter these model instances are referred to as CoVFit_Nov23_ (subscript denoting the month and year when the genotype–fitness data were obtained).

### Prediction performance of CoVFit

To evaluate the prediction performance of CoVFit_Nov23_ model instances, we examined the prediction performance of respective model instances using the corresponding test datasets. The resulting prediction performance for fitness is notably high (Spearman’s correlation: 0.992) (**Figs. 2A and 2B**). The model successfully predicted the higher fitness of later-emerging variants (**Fig. 2C**). Also, prediction of escape ability from neutralization by mAbs reaches moderately high performance (Spearman’s correlation for each epitope class: 0.551–0.810) (**Figs. 2A and S3A**). The model’s higher predictive performance was supported also in evaluations stratified by sampling country and mAb type for fitness and immune escape ability, respectively (**Fig. S3B and S3C**). Together, we show that CoVFit has sufficient power to represent the fitness landscape as well as the effect of mutations on evasion from diverse types of mAbs.

**Fig. 2.**
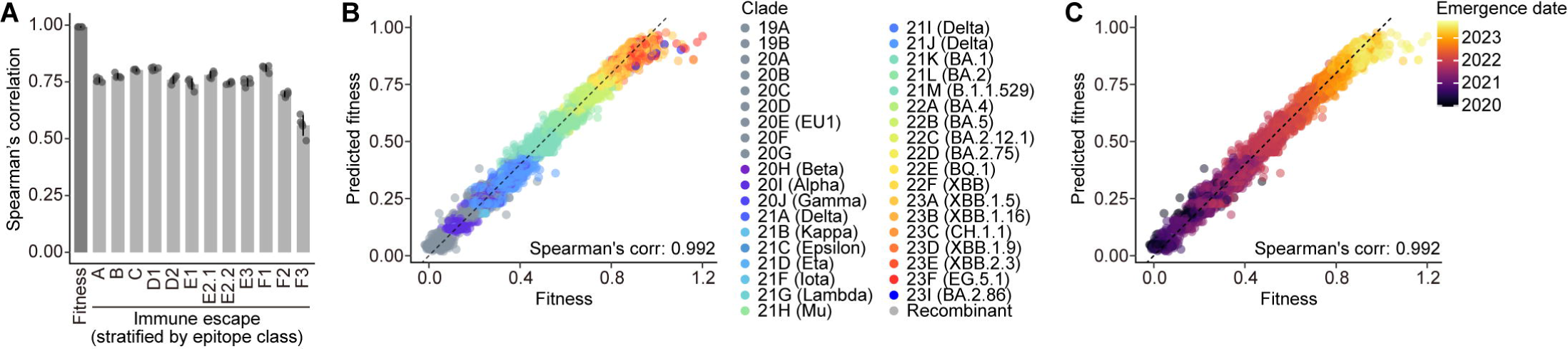
Prediction performance of CoVFit. A) Spearman’s correlation scores for predicted relative fitness values and mAb neutralization escape scores. Each cross-validation fold’s score is represented by a dot, with the mean (bar plot) and standard deviation (error bar). The correlation for mAbs was calculated in each epitope group. B) Scatter plot for fitness prediction, aggregating results from five-fold cross-validation. Dot denotes the result of a certain viral genotype in a specific country. Dot is colored by the Nextclade clade. The relative fitness value was scaled so that the 0.1 percentile and 99.9 percentile points fall between 0 and 1. A dashed line with a slope 1 and intercept 0 is shown. C) Scatter plot inherited from (**B**) but colored by the emergence date of each genotype.

### Prediction performance of CoVFit for unknown, future variants

This study aims to develop a predictive model capable of accurately assessing the fitness of yet-to-emerge variants in addition to known variants. However, the prediction performance for variants unknown to the model, particularly those that appear further in the future (after the model’s creation), cannot be evaluated if the training data contain variants highly similar to those in the test dataset, like in the performance test above (**Fig. 2**). To evaluate the model’s performance for these future variants, we synthesized datasets for ‘past’ and ‘future’ variants by splitting the existing genotype–fitness dataset based on variants’ emergence dates (**Fig. 3A**). Subsequently, we generated five instances of CoVFit solely using the past variant dataset with the five-fold cross validation scheme and then evaluated its performance on the future variant dataset.

**Fig. 3.**
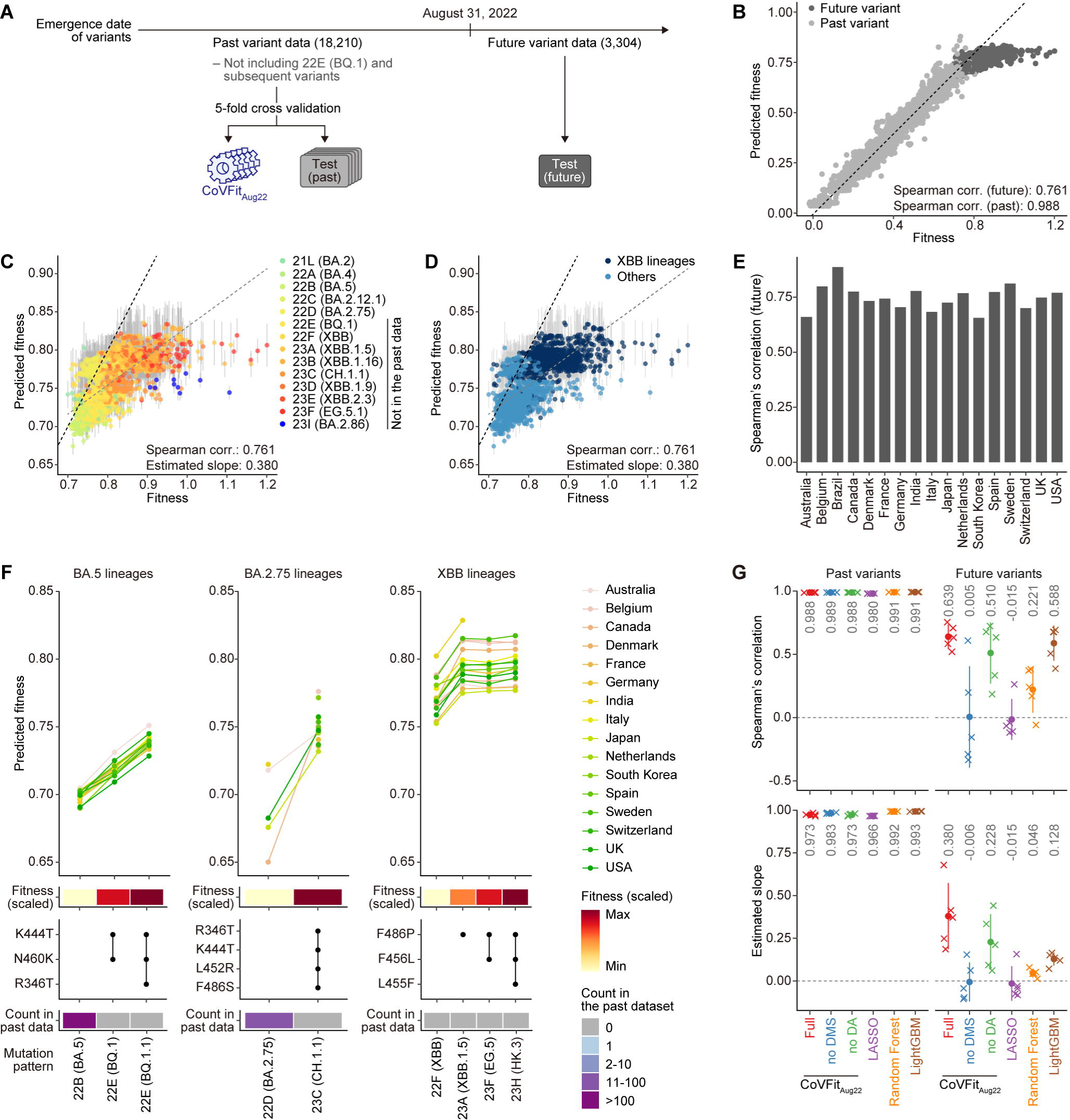
Prediction performance of CoVFit for unknown, future variants. A) Strategy to evaluate prediction performance for future variants. Model instances, referred to as CoVFit_Aug22_, were trained using data for variants emerged before August 31, 2022. The model’s prediction performance for future variants was then evaluated on data for variants emerged after this date. B) Scatter plot for fitness prediction, aggregating results from five-fold cross-validation. Both past (light gray) and future (gray) variants are included. Regarding future variants, the mean prediction across five-fold cross-validation datasets is shown. Spearman correlation for future variants was calculated using the mean prediction values. A dashed line with slope 1 and intercept 0 is shown. C) Scatter plot for fitness prediction including only future variants. Mean (dot) and standard deviation (error bar) across the five-fold prediction results are shown. Color denotes the Nextclade clade classification. In addition to the line with slope 1 and intercept 0 (black), the estimated linear regression line, based on the mean prediction values, (gray) is shown. D) Scatter plot inherited from (**C**) but colored according to whether a variant belongs to the XBB lineage. E) Spearman’s correlation scores for the fitness of future variants in each country. F) Comparison of predicted fitness among major variants. Predicted fitness value in each country (**top**), mean observed fitness value across countries (**second from top**), representative mutations defining these variants (**second from bottom**), and number of variant sequences included in the past dataset (**bottom**), are shown. G) Comparison of prediction performance among methods. Spearman’s correlation score (**top**) and estimated regression slope (**bottom**) are shown. Each cross-validation fold’s score is represented by a cross, with the mean (dot) and standard deviation (error bar). Numbers in gray denote the mean values.

The genotype–fitness dataset was thus split using a cutoff date of August 31, 2022. Applying this cutoff, Omicron lineages that emerged between late 2022 and 2023, including BQ.1 (and its sublineage BQ.1.1; clade 22E), CH.1.1 (clade 23C), XBB lineages (clades 22F, 23A, 23B, 23D, 23E, and 23F), and BA.2.86 (clade 23I), were excluded from the past datasets (**Fig. 3A**). This cutoff allows us to test the prediction performance for fitness elevation in two different evolutionary scenarios: sequential evolution (the emergence of BQ.1 and CH.1.1 from BA.5 and BA.2.75, respectively) and saltation-like evolution (the emergence of XBB from BA.2).

The trained model instances, CoVFit_Aug22_, successfully predicted the fitness of future variants to be higher than that of past variants (**Fig. 3B**). Furthermore, although CoVFit_Aug22_ tended to underestimate the fitness of future variants (estimated slope: 0.380), a higher positive correlation was still observed, even among these future variants (Spearman’s correlation: 0.761) (**Figs. 3C and 3D**). Positive correlation in future variants was observed to be robust in the analysis results stratified into each country (**Fig. 3E**). These results suggest that CoVFit_Aug22_ has the ability to predict the relative fitness of future variants. Furthermore, although the XBB lineage had 14 unique mutations in the S protein when it emerged^13^, CoVFit_Aug22_ successfully predicted the higher fitness of XBB lineages compared to other future variants including BQ.1.1, highlighting the high generalizability of CoVFit (**Fig. 3D**).

To provide a more detailed evaluation of the performance of CoVFit_Aug22_, we compared predicted fitness across major variants (**Fig. 3F)**. Within the BA.5 lineage, BQ.1 emerged from BA.5 through the acquisition of K444T and N460K substitutions in the S protein. BQ.1.1 then evolved from BQ.1, gaining an additional R346T substitution. CoVFit_Aug22_ successfully predicted the fitness elevation during sequential evolution from BA.5 to BQ.1.1. Similarly, it successfully predicted that CH.1.1, a descendant of BA.2.75 with substitutions R346T, K444T, L452R, and F486S, exhibits higher fitness than BA.2.75. Furthermore, CoVFit_Aug22_ successfully predicted that the fitness of XBB lineages is increased by the acquisition of F486P, a critical substitution in making XBB the dominant lineage^14^. This accuracy is noteworthy, considering that the past dataset did not include the XBB sequence. On the other hand, CoVFit_Aug22_ failed to accurately predict the additional increase in fitness for variants such as EG.5.1, and HK.3 compared to XBB.1.5, instead viewing them as equivalent for most countries.

We next set an additional computational experiment to evaluate the prediction performance of the model for BA.2.86, another lineage that emerged through a saltation-like evolution from BA.2, marked by the acquisition of approximately 30 mutations in the S protein^17^ (**Fig. S4)**. In this analysis, the fitness dataset was split according to the cutoff date of July 31, 2023, to exclude data for BA.2.86 lineages (including its descendant lineage, JN.1) from the past dataset (**Fig. S4)**. We then generated additional model instances, CoVFit_Jul23,_ using this past dataset. Upon evaluation with the respective future dataset, CoVFit_Jul23_ successfully predicted the higher fitness of BA.2.86 over BA.2—the parental lineage of BA.2.86—and over other lineages including BA.5, BQ.1, and XBB (**Fig. S4B)**. However, it failed to predict the higher fitness of BA.2.86 over XBB descendant lineages, such as XBB.1.5, EG.5, and HK.3. Our analyses demonstrate the high generalizability of CoVFit as well as provide insight into its limitations, which taken together allow the responsible interpretation of its results for making specific predictions on variant fitness.

### Performance comparison per-component and other prediction models

To assess the impact of incorporating DMS data into CoVFit on the efficacy of fitness prediction, we generated an additional model instance, CoVFit_noDMS_, by training without the DMS dataset and subsequently evaluated its predictive performance against the original model (**Figs. 3G and S5**). This reduced model was assessed through experiments employing a past–future variant data splitting method, with a cutoff date of August 31, 2022. In predicting the fitness of past variants, CoVFit_noDMS_ exhibited prediction performance similar to the original CoVFit in terms of both Spearman’s correlation and the estimated regression slope. However, CoVFit_noDMS_ substantially underperformed in predicting the fitness of future variants. Similarly, we examined the impact of the domain adaptation step on prediction performance and demonstrated the contribution of this step in achieving higher performance (**Figs. 3G and S5**). Removing the DMS dataset had a much stronger effect than omitting the domain adaptation step, underscoring the critical role of DMS dataset incorporation. Collectively, CoVFit’s superior predictive performance can be attributed to its incorporation of functional information on mutations and the application of domain adaptation frameworks.

Next, to assess the performance of CoVFit in comparison with other methods, we constructed alternative prediction models using non-deep learning methods, such as LASSO, Random Forest, and Light Gradient Boosting Machine (LightGBM) (**Figs. 3G and S5**). In predicting the fitness of past variants, these models exhibited prediction performance similar to CoVFit in terms of both Spearman’s correlation and the estimated regression slope. However, when predicting the fitness of future variants, CoVFit surpassed both the linear model (LASSO) and the more complex decision tree-based models (Random Forest and LightGBM). Particularly, the regression slope estimate of CoVFit was substantially higher than those of other models, indicating a smaller underestimation bias in CoVFit than other methods.

### Fitness elevation events during SARS-CoV-2 evolution

To deepen our understanding of the fitness landscape of SARS-CoV-2, we developed a CoVFit-based phylogenetic framework to analyze fitness elevation throughout its evolution (**Fig. 4)**. First, we constructed a phylogenetic tree of 11,098 variants, which correspond to viral genome sequences encoding respective S protein genotypes. Subsequently, ancestral S protein sequences at the internal nodes of the tree were reconstructed (**Fig. 4A)**. We then inferred the fitness of all nodes, representing both observed and reconstructed ancestral sequences, utilizing the latest CoVFit_Nov23_ models (**Figs. 4B and S6** for Omicron and all lineages, respectively). We obtained five predicted fitness values per node with according mean and standard error using model instances generated via a five-fold cross-validation scheme. Finally, we identified branches where fitness elevation was statistically significant (false discovery rate; FDR<0.1) by comparing predicted fitness values between a given node and its parent node. Of the 9,846 branches that acquired mutations in the S protein, 549 (5.6%) branches were identified with significant fitness elevation (**Fig. S6**), including 334 branches within the Omicron lineages (**Fig. 4C**). We observed increases in viral fitness both in the branches representing the most recent common ancestor (MRCA) of major lineages and throughout their subsequent diversification (**Figs. 4C and S6)**.

**Fig. 4.**
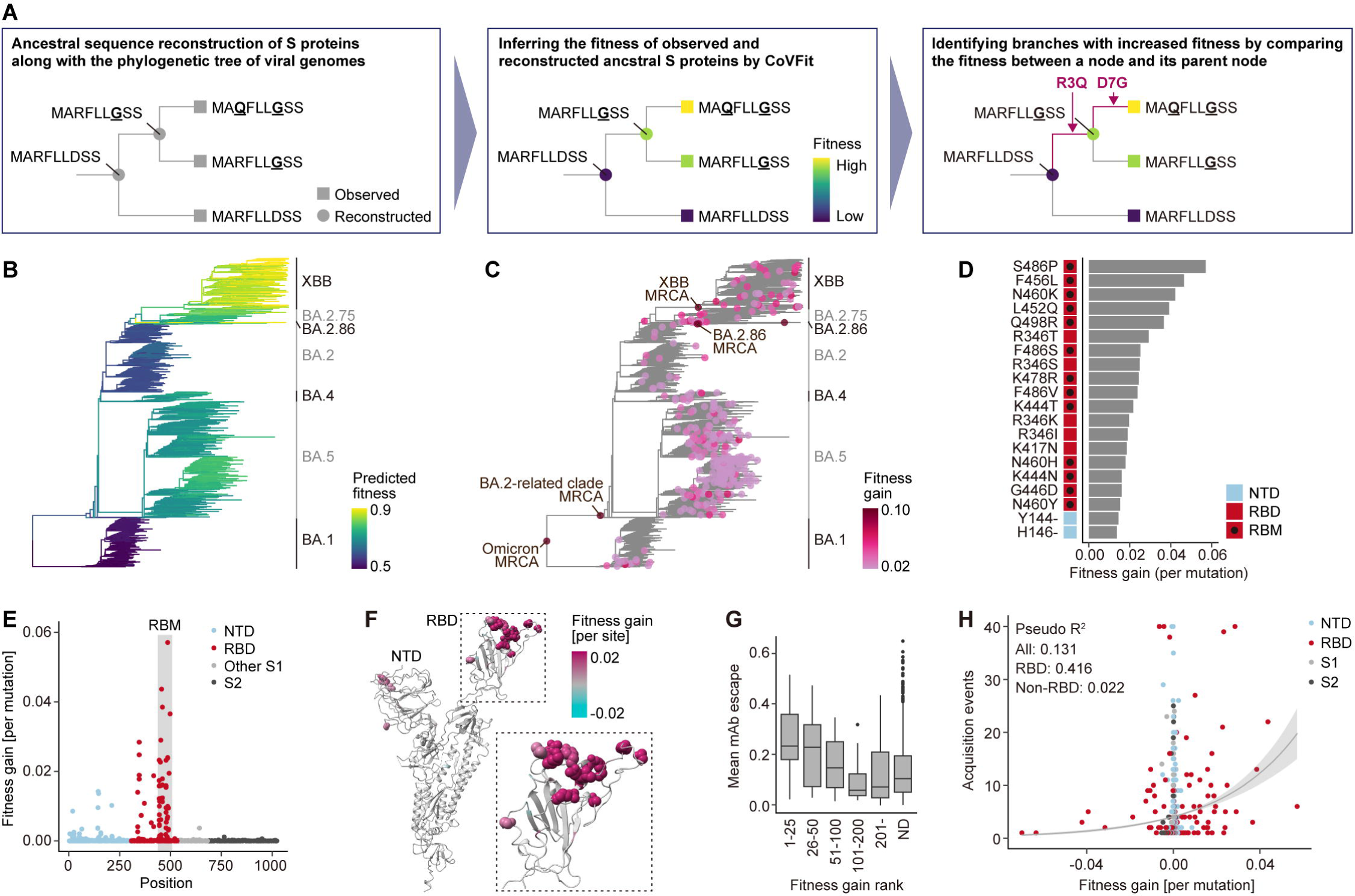
Detection of fitness elevation events during Omicron diversification. A) Scheme to detect phylogenetic branches with fitness elevation utilizing CoVFit models. B) Inference of change in fitness through Omicron’s evolution. The maximum likelihood (ML) tree of Omicron lineages is shown. Branch color indicates an inferred fitness value for each phylogenetic node, including both observed and reconstructed ancestral genotypes of S proteins in the phylogenetic tree. C) Detection of fitness elevation events during Omicron’s evolution. Dot color indicates inferred fitness gain in each branch, calculated as difference in predicted fitness between a node and its parental node. D) Mean fitness gain over a specific mutation during Omicron evolution. Since some mutations have been acquired multiple times, the mean value of fitness gain among acquisition events was used as the “fitness gain [per mutation]” score. Top 20 mutations regarding this score are shown with the protein domain information. E) Enrichment of fitness-associated mutations in the RBD, particularly in its RBM. The negative score is clipped to 0. F) Mapping the site-wise fitness gain score on the 3D structure of the ancestral D614G S protein (PDB: 7BNN)^49^. If multiple mutation types are present in a specific site, the maximum value is shown as the “fitness gain [per site]” score. Amino acid side chains for the top 15 sites regarding this score are shown as sphere. The plot was generated using Chimera X^50^. G) Association of fitness gain rank with the mean mAb escape score. This escape score was calculated as the mean of the escape score across mAbs over a mutation. The ND group includes mutations not observed in our phylogenetic analysis. H) Association of the fitness gain [per mutation] score with the inferred acquisition count. Estimated regression curve (line) with standard error (ribbon) by Poisson regression using all mutations are shown. In addition, the Nagelkerke’s pseudo R^2^ values for Poisson regression analyses using all mutations, RBD mutations, and non-RBD mutations are shown.

To identify mutations critical for fitness elevation, we next calculated the average fitness gain per mutation by examining branches with the acquisition of a single mutation (**Figs. 4D and S6** for Omicron and all lineages, respectively). Mutations with higher fitness gain are predominantly found in the RBD of the S protein, particularly in its receptor binding motif (RBM), consisting of the binding interface to ACE2^30^ (**Figs. 4E, 4F, and S6)**. Additionally, we identified significant non-RBD mutations, like the Y144- and H146-deletions, in the N-terminal domain. We found that mutations with a greater impact on fitness elevation also tend to enhance the virus’s ability to evade humoral immunity (**Fig. 4G)**. Furthermore, mutations with significant fitness impact in Omicron lineages tend to have been acquired multiple times in a convergent manner throughout Omicron’s evolution (**Fig. 4H)**. This association was evident in RBD mutations (Nagelkerke’s pseudo R^2^: 0.416) but not in non-RBD mutations (Nagelkerke’s pseudo R^2^: 0.022).

### Context-specific effects of the F456L substitution

We found that acquisitions of some mutations were overrepresented in a specific phylogenetic lineage. For example, while the R346T substitution was convergently acquired across Omicron lineages, the F456L substitution was markedly overrepresented in the XBB lineage (**Fig. 5A)**. To quantify differences in the fitness effect of F456L among lineages, we performed *in silico* mutational scanning analysis leveraging CoVFit_Nov23_ by computationally inducing F456L in various S protein backbones and inferring the fitness gain caused by this substitution (**Fig. 5B**). Predicted fitness gain by F456L in the XBB lineage was clearly higher than those in other lineages. Together, these results suggest that the fitness-increasing impact of F456L is specific to the XBB lineage.

**Fig. 5.**
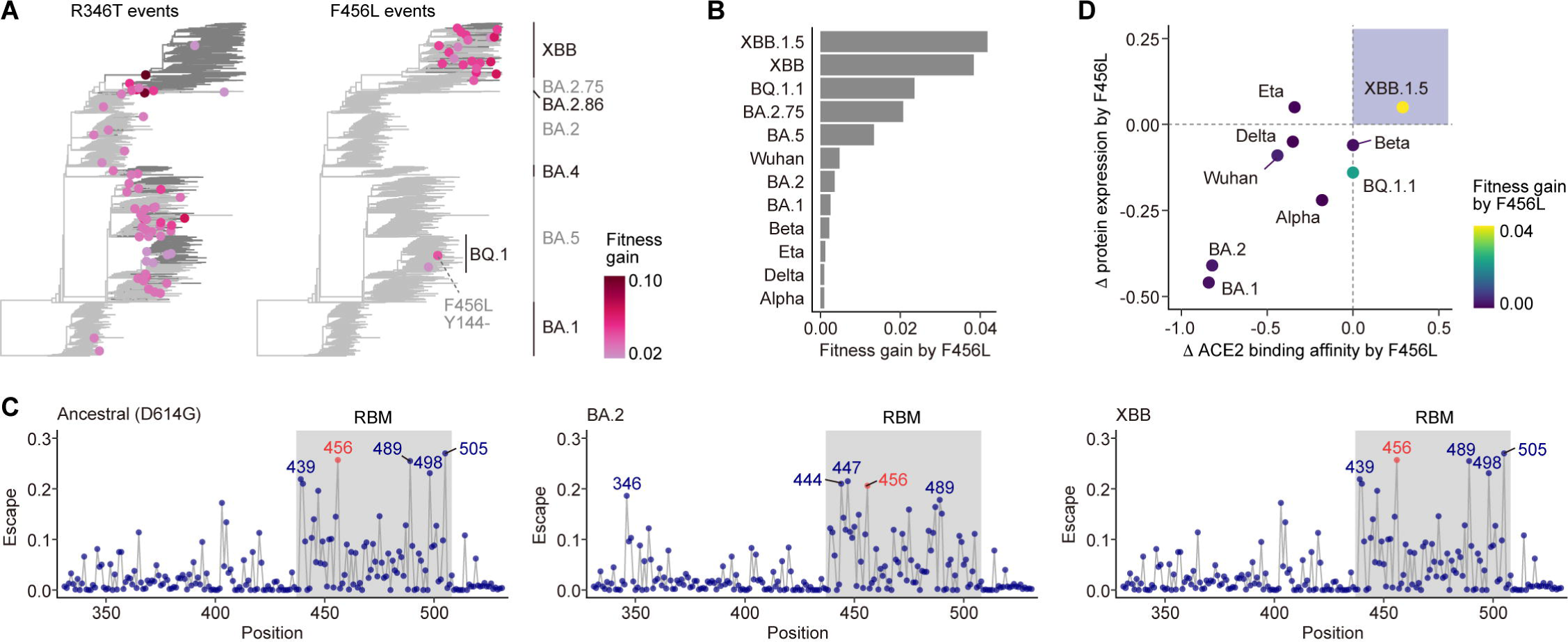
Context specific effect of the F456L substitution. A) Examples of convergent acquisitions of specific substitutions. A node indicates the acquisition events, and node color denotes fitness gain at the acquisition events. Branch color denotes the presence (gray) or absence (light gray) of specific substitutions in the reconstructed ancestral S protein sequences. B) Fitness gain upon F456L in each backbone S protein sequence, inferred by *in silico* mutational scanning using CoVFit. C) Site-wise immune escape score for the ancestral D614G strain, BA.2, and XBB variants, estimated by mAb escape estimator^32^ based on Cao’s DMS data^29^. Top 5 sites regarding the escape score are annotated. D) Effect of F456L on the S protein’s expression (stability) and ACE2-binding affinity, extracted from publicly available DMS data from Taylor and Starr^31^. Dot color indicates inferred fitness gain shown in (**B**). Higher values indicate enhanced higher expression and ACE2 binding affinity values.

To gain mechanical insights into the XBB-specific F456L effect on viral fitness, we analyzed published DMS data. This includes the data on mAb neutralization by Cao et al.^29^ and those on the RBD ACE2-binding affinity and protein stability by Taylor and Starr^31^. Substitutions at F456 has one of the largest impacts on neutralization escape, according to the escape estimator data^32^, in various lineages including the ancestral D614G strain’s S protein (**Fig. 5C)**. On the other hand, the effects of this mutation on ACE2 binding and protein expression were different among S protein backbones. While F456L enhances ACE2 binding and protein expression in the XBB S protein, this substitution has a negative effect on ACE2 binding and/or protein expression in all tested S backgrounds other than XBB (**Fig. 5D)**. The non-deleterious effect of F456L in ACE2 binding and expression, unique to XBB, has been confirmed in both RBD and full S DMS assays in previous studies^29,33^. These results suggest that F456L confers preferable effects on the XBB’s S protein but confers a double-edged sword effect on other lineages’ S protein. Together, the XBB-specific positive effect of F456L on fitness can be explained by the XBB-specific removal of deleterious effects of this mutation. This example validates the effectiveness of CoVFit to infer mutational effects in a context-specific manner.

### CoVFit-based *in silico* DMS to predict the fitness of possible variants

To demonstrate the utility of CoVFit in predicting the emergence of variants with increased fitness, we developed a simulation method, namely CoVFit-based *in silico* DMS. In this method, we computationally introduced every possible single amino acid substitution into the S protein of JN.1 (BA.2.86.1.1)—the dominant variant as of March 2024. Subsequently, the fitness gain from each substitution was inferred using CoVFit_Nov23_. It is important to note that the training dataset for CoVFit_Nov23_ includes only a single sequence of JN.1 with no sequences of its descendant lineages.

This *in silico* DMS analysis identified substitutions at residues F456, K478, and R346 as having the highest predicted fitness gains (**Figs. 6A and 6B**). To validate these predictions, we compared them with the latest genome surveillance data as of March 11, 2024. Notably, emerging JN.1 sublineages carrying substitutions at F456, K478, R346, or the combination of F456 and R346 are rapidly expanding within the global JN.1 population (**Fig. 6C**). These results indicate that CoVFit_Nov23_ successfully predicted the higher fitness of these JN.1 sublineages compared with the parent JN.1—effectively before their emergence or detection. Together, these results highlight the capability of CoVFit-based simulation analysis to predict in advance the emergence of variants with a higher potential for epidemic spread.

**Fig. 6.**
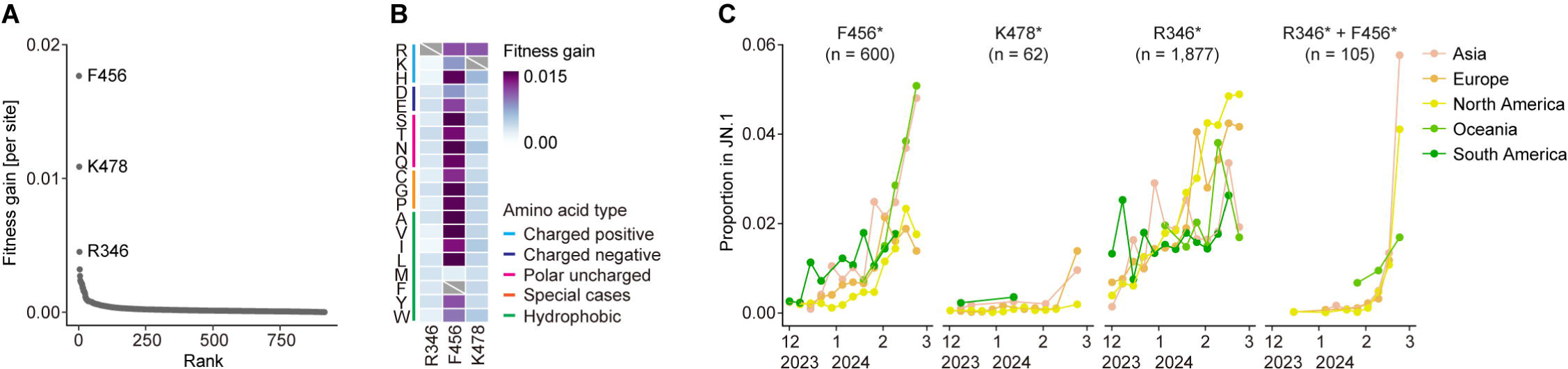
CoVFit-based *in silico* DMS on the JN.1 variant. A) Ranking of amino acid sites regarding the fitness gain [per site] score inferred by *in silico* DMS with CoVFit_Nov23_ on the S protein of JN.1. Only sites with positive score are shown. B) Heatmap depicting inferred fitness gain by each amino acid substitution. C) Change in frequency of JN.1 subvariants harboring substitutions in specific amino acid sites, within the JN.1 population. The genome surveillance data from December 1, 2023, to February 29, 2024, was used. Frequencies were calculated using 7-day bins. Total sequence number is shown for each viral group.

### CoVFit-CLI tool

The trained CoVFit_Nov23_ model instances are available as a command line tool, CoVFit-CLI, from our GitHub repository (https://github.com/TheSatoLab/CoVFit). Researchers can use the stand-alone program to conveniently provide fitness and DMS escape predictions for their own SARS-CoV-2 S protein sequences. The CoVFit-CLI tool will receive periodic updates to the model instances, trained on the latest genomic surveillance data.

## Discussion

In this study, we established CoVFit, a protein language model to predict the relative fitness of SARS-CoV-2 variants with high generalizability. Previous studies have proposed machine learning models predicting viral phenotypes strongly associated with fitness, such as immune evasion ability^34,35^ or protein *grammaticality*^36,37^. However, to our knowledge, CoVFit is the first machine learning model designed to directly predict viral fitness (i.e., R_e_), which represents a variant’s ability to spread in the host population. Furthermore, we demonstrated that CoVFit can be used to understand the fitness landscape of SARS-CoV-2 and to flag variants with higher potential to spread before or immediately after their detection.

CoVFit model instances (CoVFit_Sep22_) successfully predicted the fitness increases resulting from sequential evolution, specifically from BA.5 to BQ.1.1 and from BA.2.75 to CH.1.1. Furthermore, the model instances successfully predicted the fitness increase resulting from the saltation-like evolution that led to the emergence of the XBB lineage (**Fig. 3**). Moreover, the model instances successfully predicted the first step of sequential fitness elevation in the XBB lineage at the acquisition of the F486P mutation. This robust generalizability is notable since 14 mutations were acquired in the S protein through recombination during the emergence of XBB^13^. This ability is primally acquired through the multitask learning of knowledge on variant fitness and mutational effects on immune evasion ability (**Fig. 3G**). On the other hand, we also showed that the current model does not accurately generalize to all situations. For example, CoVFit model instances (CoVFit_Sep22_ and CoVFit_Jul23_) underestimated the fitness elevation in the emergence of BA.2.86, where approximately 30 mutations were acquired at once^17^ (**Fig. S4**). Despite this underestimation, however, it is worth noting that the model did successfully predict the substantially higher fitness of BA.2.86 compared to its ancestral strain BA.2 and following successively emerged lineages like BA.5, BQ.1.1 and XBB.

CoVFit has the potential to transform the genome surveillance of SARS-CoV-2 variants with higher epidemic risk. With the prolongation of the SARS-CoV-2 epidemic and the current decline in genome surveillance efforts (https://gisaid.org/hcov-19-variants-dashboard/), methods that can directly predict the fitness of emerging variants from their genotypes are becoming increasingly critical. By employing CoVFit to scan viral genomes newly registered in a viral genome surveillance database, it is anticipated that we can identify variants likely to have higher fitness at an unprecedentedly early stage, ideally upon the registration of a single sequence. This method allows for the direct detection of viruses with higher fitness, bypassing the steps of identifying and naming new variants, which often require manual designation on a public forum (https://github.com/cov-lineages/pango-designation). This efficiency enables the continuation of variant monitoring with less time and fewer human efforts. Moreover, the use of CoVFit for *in silico* DMS has the potential to predict variants with high fitness before their actual emergence or detection, as exemplified by the successful prediction of JN.1 sublineages with enhanced growth advantage (**Fig. 6**). Given the efficient training process of CoVFit, which completes within approximately 24 hours per model instance utilizing a single Nvidia A100 GPU, the CoVFit model can be updated regularly to incorporate information on the latest variants (see **Methods**). Together, a highly efficient monitoring system for emerging variants implementing the latest CoVFit instances can be developed in future work.

In this study, we developed a framework that combines CoVFit with a phylogenetic approach to identify mutations that enhance viral fitness (**Figs. 4 and 5**). Despite the common challenge of interpretability in machine learning models, our framework is designed to assess not just the average effect of specific mutations but also their context-specific or epistatic effects. For instance, it can distinguish the varying impacts of a particular mutation (e.g., F456L) across different contexts (e.g., within the XBB lineage versus other lineages) (**Fig. 5**). This framework serves as a valuable complement to *in silico* DMS. Compared to *in silico* DMS, this approach aims to retrospectively interpret observed evolutionary steps and identify patterns in their occurrence.

Previous studies have reported that some mutations are convergently acquired through the SARS-CoV-2 evolution probably because these convergent mutations confer positive effect on viral fitness^29,38^. Indeed, some studies have inferred the importance of mutations for viral fitness according to how often these mutations are acquired convergently, under the assumption that these two features are correlated^29,39^. However, despite our prior investigations focusing on a limited set of mutations^12^, the degree of association between a mutation’s acquisition frequency and its impact on fitness has remained unclear. In this study, we comprehensively investigated this issue and found an association between these two features, with a particularly strong correlation observed in mutations within the RBD (**Fig. 4H**). This result suggests, for accurate predictions of viral evolution, it is effective to consider both the acquisition frequency of mutations and their fitness effects. By steering future development towards a machine learning model based on a phylodynamics approach integrating both factors, we could potentially achieve precise predictions and simulations of viral evolution.

Despite the utility of CoVFit, it still has several limitations in its present form. First, as the objective variable, we utilized the fitness (R_e_) of a variant, which was estimated from viral genome surveillance data and inherently carries a degree of uncertainty and bias. This bias arises partly because the surveillance does not employ random sampling. Often, samples from specific infection clusters are disproportionately represented in the surveillance data, potentially leading to a biased estimation of the fitness of certain variants. Second, the logistic model used to estimate variant fitness operates under a strong assumption that the relative fitness among variants remains constant over time^4–7^. Although this model is widely used, this assumption might not always hold true. In reality, the relative fitness of variants can vary in response to changes in the host population’s immune status, influenced by factors such as natural infections and vaccinations^2,3^. Third, in our fitness prediction task, we utilized sequences from real-world variants as training inputs. Mutations with a substantial negative impact on fitness must be under-represented in our training dataset since such mutations are usually eliminated through natural selection. Consequently, CoVFit is likely to underestimate the negative effects of certain mutations on fitness. Furthermore, it should be noted that the current model tends to underestimate the fitness of future variants (**Fig. 3**). Other technical limitations and their potential solutions are discussed in the **Room for improvement of CoVFit** section in **Methods**. Together, the fitness predictions made by our model should be carefully interpreted in the context of complimentary information.

Despite the limitations mentioned, CoVFit holds the potential to decipher the fitness landscape of viruses. Our approach is poised to contribute to the development of innovative methods for the early prediction of future epidemic variants and for advancing viral evolutionary predictions. These advancements are critical for efficient epidemic control, vaccine development, and drug discovery. Furthermore, the methodologies employed in CoVFit can be applied to predicting the fitness of other viruses including viral pathogens causing future pandemics. In anticipation of the next pandemic, it is imperative to continually develop foundational bioinformatics methods that assist in epidemic control, leveraging the extensive genomic data efforts of SARS-CoV-2 as an archetypical forerunner.

## Methods

### Preparation of genotype–fitness dataset

We retrieved all SARS-CoV-2 genome sequences and their associated metadata available as of November 2, 2023, from GISAID (https://gisaid.org/). We then assigned the most recent PANGO lineage classification available at that time to each sequence in our dataset using Nextclade v.2.14.0^40^ with Nextclade dataset version “2023-10-26T12:00:00Z”. Then we excluded low quality sequences based on the following criteria: (i) absence of collection date information; (ii) samples derived from animals other than humans; or (iii) more than 1% undetermined nucleotide characters.

To develop a classification system for SARS-CoV-2 with a higher resolution than the PANGO lineage and based solely on the sequence of the S protein, we defined the genotype of the S protein and utilized it as the virus classification system in this study. The S protein genotypes refer to groups of viral sequences that share a unique set of mutations in the S protein. To achieve this, we first identified mutations in the S protein observed in more than 100 sequences. We then analyzed the mutation patterns across each sequence, enabling us to categorize sequences into genotypes based on these patterns. Only genotypes represented by 20 or more sequences in any country were considered for our analysis. As a result, a total of 13,643 S protein genotypes were included in our dataset. Each genotype was linked to the Nextclade PANGO lineage, Nextclade clade, and a representative genome sequence. This representative sequence was randomly chosen from the collection of sequences of the genotype. The emergence date of a genotype was determined as the 1st percentile date of collection for the viral isolates within the genotype. The viral genome sequences contained within this dataset are summarized under the EPI_SET_ID: EPI_SET_240311ma, which can be accessed through the GISAID website (https://gisaid.org/).

To estimate the relative R_e_ value of each genotype in each country, we began by tallying the daily count of each genotype within each country’s dataset. We applied a multinomial logistic model to the count data of each country and estimated parameters using the maximum likelihood method with the ‘multinom’ function in the ‘nnet’ package v.7.3.18 in R v.4.2.1. We then extracted the estimated growth rate (slope parameter) of each genotype relative to a reference genotype. The reference genotype was chosen as the one with the highest detection count in the country with the lowest overall detections. Accordingly, the major genotype of the BA.5 lineage (equivalent to BA.5.2.1 in the PANGO lineage) was selected as the reference. The relative R_e_ of each viral lineage, *r_l_*, was calculated according to the slope parameter β*_l_* as *r_l_* = exp(γβ*_l_*), where β*_l_* and *r_l_* are the logistic slope parameter and relative R_e_ of viral lineage l, respectively, and γ is the average viral generation time (2.1 days) (http://sonorouschocolate.com/covid19/index.php?title=Estimating_Generation_Time_Of_Omicron)^8^. This model has two strong assumptions that relative fitness among variants remains constant over time and that all variants have the same the average viral generation time^4–7^ (**See Discussion**). Estimated R_e_ value for each genotype is summarized in **Table S1**.

Prior to the training step, we excluded S protein sequences containing more than 5 ambiguous characters and more than 30 amino acid deletions from the dataset. Also, we focused on countries where more than 300 genotypes were detected, which led to retaining data for 17 countries: Australia, Belgium, Brazil, Canada, Denmark, France, Germany, India, Italy, Japan, Netherlands, South Korea, Spain, Sweden, Switzerland, UK, and USA. As a result of this additional filtering, a total of 21,571 genotype–fitness (relative R_e_) data points, encompassing 12,914 genotypes across 17 countries, were included in our genotype–fitness dataset.

The estimated fitness value was transformed using the natural logarithm function, and then the data was scaled so that the 0.1 percentile and 99.9 percentile points fall between 0 and 1 before model training.

### Preparation of DMS data for mAbs evasion

In this study, we utilized DMS data on evasion from mAbs provided by Cao et al.^29^. The processed DMS data, specifically the mutation-wise immune escape score prepared for the antibody-escape estimator developed by Greaney et al.^32^, was downloaded from the Bloom lab GitHub repository on April 11, 2023 (https://github.com/jbloomlab/SARS2_RBD_Ab_escape_maps/blob/main/processed_data/escape_data_mutation.csv). We applied specific exclusion criteria to the DMS data: i) mAbs categorized as “SARS convalescents” and “WT-engineered”; and ii) mAbs with an IC_50_ value ≥10, indicative of very weak binding affinity, for the target virus. The escape score in this repository was calculated using a DMS experiment using the ancestral D614G strain’s RBD. Following the methods of Greaney et al., we defined a weighted escape score for each target virus (e.g., D614G and BA.2) from this escape score, following the method of Greaney et al. Specifically, the escape score was multiplied by the IC_50_ value for the S protein of the target virus, followed by negative log transformation with pseudo count 1. The weighted escape score was scaled so that 0 and 95 percentile fell within the range 0–1, and values above 95 percentile were clipped to 1. For the comprehensive training of CoVFit, the weighted escape values for D614G were employed. On the other hand, considering that variants predominantly circulating after early 2022 are related to the BA.2 lineages, the weighted escape values for BA.2 were used in the training for CoVFit_Aug22_ and CoVFit_Sep23_.

### Dataset preparation for domain adaptation

The S protein sequences for *Coronaviridae*, except for SARS-CoV-2, were downloaded from the NCBI Identical Protein Groups database (https://www.ncbi.nlm.nih.gov/ipg) on July 3, 2023, using the following search query: query:(“Alphacoronavirus”[Organism] OR “Betacoronavirus”[Organism] OR “Gammacoronavirus”[Organism] OR coronavirus[All Fields]) AND (spike[All Fields] OR S[All Fields] OR surface[All Fields]) NOT (“Severe acute respiratory syndrome coronavirus 2”[Organism] OR (“Severe acute respiratory syndrome coronavirus 2“[Organism] OR (“Severe acute respiratory syndrome coronavirus 2”[Organism] OR (“Severe acute respiratory syndrome coronavirus 2“[Organism] OR (“Severe acute respiratory syndrome coronavirus 2”[Organism] OR (“Severe acute respiratory syndrome coronavirus 2“[Organism] OR (“Severe acute respiratory syndrome coronavirus 2”[Organism] OR (“Severe acute respiratory syndrome coronavirus 2“[Organism] OR (“Severe acute respiratory syndrome coronavirus 2”[Organism] OR (“Severe acute respiratory syndrome coronavirus 2“[Organism] OR SARS-CoV-2[All Fields])))))))))) NOT (“unidentified”[Organism] OR (“unidentified”[Organism] OR (“unidentified”[Organism] OR (“unidentified”[Organism] OR (“unidentified”[Organism] OR “Unknown”[All Fields]))))) NOT “unidentified human coronavirus”[Organism] NOT (“synthetic construct”[Organism]) AND (“1000”[SLEN]: “1500“[SLEN]). The metadata associated with these sequences were also downloaded. The S protein for SARS-CoV-2 Wuhan-Hu-1 strain was downloaded using NCBI Datasets Command-line tools v.15.6.1 (https://www.ncbi.nlm.nih.gov/datasets/docs/v2/download-and-install/) and subsequently incorporated into our dataset. Sequences with more than 5% unidentified amino acids were filtered out. Next, we removed redundant sequences using CD-HIT v.4.8.1^41^ with a clustering threshold of 99% sequence identity. However, even after the CD-HIT filtering, a large number of sequences (i.e., 796 sequences) corresponding to the porcine epidemic diarrhea virus (PEDV) remained in the dataset. To reduce the redundancy of this virus group, we randomly selected 10 representative PEDV sequences to retain in our dataset. Consequently, 1,392 *Coronaviridae* sequences were included in our dataset.

In addition to the *Coronaviridae* S protein dataset, we prepared a dataset for the SARS-CoV-2 S protein specifically for domain adaptation. Of the S protein genotypes we defined in this study (see the section **Preparation of genotype– fitness dataset**), we eliminated genotypes with an emergence date later than August 31, 2022, in order to prevent the model from accessing data beyond this cutoff date during the domain adaptation process. Subsequently, we removed redundant sequences using CD-HIT with a clustering threshold of 99%, in accordance with the method described above. Consequently, 114 S protein sequences of SARS-CoV-2 were retained in the dataset. Finally, we combined the *Coronaviridae* and SARS-CoV-2 S protein datasets, resulting in 1,506 sequences, for use in domain adaptation. The S protein sequence information used in the domain adaptation step is summarized in **Table S2**.

### Introduction of CoVFit

We developed CoVFit, a fitness prediction model based on S protein sequences, by fine-tuning the ESM-2 protein language model (**Fig. 1A**). To enhance the model’s performance, we employed three key techniques: domain adaptation, multitask learning, and Low Rank Adaptation (LoRA)^42^. Domain adaptation is an additional pretraining phase using a custom data collection. In this study, we performed domain adaptation using S protein sequences from various human and animal coronaviruses (see the **Dataset preparation for domain adaptation** section). This technique enabled the model to better learn the general properties of these proteins (see the **Domain adaptation** section). Multitask learning, a framework that trains a model on different types of data simultaneously, was utilized to allow the model to capture critical information shared across tasks, thereby enhancing its generalization capabilities. In CoVFit, the model was finetuned using a total of 1,565 regression tasks including to predict fitness values for 17 countries and to predict relative binding affinity for 1,548 mAbs (see the **Model architecture of CoVFit** section). Finally, LoRA is a technique to fine-tune large models efficiently that reduces the requirements for GPU memory resources without compromising the model’s prediction performance. LoRA additionally contributes to mitigating the model’s tendency to overfit (see the **Model finetuning and performance evaluation** section for further information).

The input for CoVFit consists of amino acid sequences of SARS-CoV-2 S proteins, aligned with the S protein of the Wuhan-Hu-1 strain. These aligned sequences can be generated by Nextclade. CoVFit can predict the fitness (R_e_) value across 17 countries and the ability to evade 1,548 types of mAbs for a given S protein sequence (**Fig. S1A**).

Training of the CoVFit model completes within 24 hours on a computational node with a single Nvidia A100 GPU (40GB) for each instance. Consequently, the model can be updated routinely using the latest genome surveillance data without intensive computational resource requirement.

Utilizing a five-fold cross-validation scheme, we generated five instances of the CoVFit model, which enabled us to estimate both the average prediction value and its uncertainty across these models (**Fig. 1B**). This approach was chosen because the predicted values, especially regarding the fitness of future variants, can vary among different instances of the trained models (**Fig. 3**).

### CoVFit implementation

ESM-2 models with various parameter sizes are available^27^ (https://github.com/facebookresearch/esm). Of these models, we used the version with 650M parameters, prioritizing a balance between prediction performance and computational cost. According to the official benchmark using the unsupervised contact prediction task, this 650M parameter model achieves a performance 1.7 times superior compared to the 35M parameter model, which possesses 20 times fewer parameters. However, the performance gain when comparing the 650M model to the 15 billion (B) parameter model, which has 20 times more parameters, is relatively modest at only 1.08 times. Furthermore, a systematic analysis presented in a recent preprint indicates that enlarging the parameter size of a protein language model does not necessarily enhance prediction performance for tasks outside of protein structure prediction^43^. Given this insight, we opted not to employ models larger than the 650M model, such as the 3B or 15B models, in our study.

The ESM-2 model has a maximum input sequence length (1,024 amino acids) due to the computational demands of self-attention, which requires memory in proportion to the square of the sequence length (O(L^2^)). Unfortunately, the S protein of the Wuhan-Hu-1 strain is composed of 1,273 amino acids, exceeding the model’s limit. Consequently, amino acid sequences beyond the 1024th position (amino acids 1,025–1,273; the C-terminus of the S2 subunit) are truncated and not utilized in the ESM-2 model. This constitutes a technical limitation of CoVFit. Nonetheless, this limitation is anticipated to minimally impact performance, considering that while mutations predominantly occur within the S1 subunit (amino acids 1–681), the S2 subunit (amino acids 682–1,273) remains highly conserved and with fewer mutations.

CoVFit was implemented using Python v.3.11.4, NVIDIA CUDA v.12.1.0, PyTorch v.2.1.0, Transformers v.4.31.0, and PEFT v.0.5.0. Further information about the system requirements for CoVFit can be found in the GitHub repository (https://github.com/TheSatoLab/CoVFit). The computational codes were executed on a supercomputer node equipped with a single NVIDIA A100 GPU with 40GB RAM unless otherwise noted.

### Domain adaptation

To establish the ESM-2_Coronaviridae_ model, domain adaptation was carried out using the masked language learning scheme as described in Delvin et al.^44^. For domain adaptation, the S protein dataset prepared in the **Dataset preparation for domain adaptation** section was used. For our model’s domain adaptation training, each input sequence had 15% of its positions masked randomly, with each instance of a position’s masking having an 80% chance to be a <MASK> token, a 10% chance to be incorrect, and a 10% chance to be the original. Subsequently, amino acid or token types for these 15% of positions were predicted in batched training steps and model weights were updated using a cross-entropy loss function.

Using the scheme above, we trained the 650M parameter ESM2 model with provided MaskedLM layer, downloaded via functions implemented in the Hugging Face Transformers library. The model was trained for 30 epochs. Batch size was set at 5. A base learning rate of 2e−5 was used with one epoch of warmup, and a cosine-based learning rate scheduler was implemented to successively lower the learning rate during training.

To compare the inference ability of the ESM-2_Coronaviridae_ model to the original ESM-2 model, we performed inference with both models on masked SARS-CoV-2 S protein sequences. Since our dataset for domain adaptation training includes the S proteins of genotypes emerged up to August 31, 2022, we used genotypes with emergence dates later than September 1, 2022, for inference. The same masking parameters as in the training were used. The results on the test dataset were converted to perplexity scores as the exponential of the cross-entropy loss value calculated during inference. Given as *perplexity* = e^−∑*xP*(*x*)log*Q*(*x*)^ where *P*(*x*) is the true probability distribution and *Q*(*x*) is the probability distribution from the model’s predictions, the perplexity score represents how certain the model is in making its predictions, with lower values demonstrating higher certainty. For our inference results, the original ESM-2 model produced a perplexity score of 11.38, whereas the ESM-2_Coronaviridae_ model achieved a low perplexity score of 1.17, demonstrating higher prediction certainty after domain adaptation training (**Fig. S1C**).

To assess the possibility of the domain adaptation negatively impacting the model’s original ability to provide inference on a wide variety of proteins, we again compared the original EMS-2 and ESM-2_Coronaviridae_ models, this time with protein sequences sampled from the UniRef50 released in March 2018 (https://dl.fbaipublicfiles.com/fair-esm/pretraining-data/uniref201803_ur50_valid_headers.txt.gz). A subset consisting of 29,950 sequences were randomly sampled from the full 3,016,211 sequences and used for the evaluation. The perplexity values of the models were checked as above for the two models, with the original ESM-2 model’s perplexity score at 6.76 and the ESM-2_Coronaviridae_ model’s perplexity score at 6.86, demonstrating that the model retains its certainty on general proteins after domain adaptation (**Fig. S1C**).

We conducted the domain adaptation training in Python v3.10.9 with CUDA v12.1.1 and torch v2.1.0.dev20230601 using the Hugging Face Transformers v4.34.1 library.

The computation was executed on a single NVIDIA RTX 6000 Ada GPU with 48GB RAM. More detailed information on implementation is available in the GitHub repository (https://github.com/TheSatoLab/CoVFit).

### Model architecture of CoVFit

For the multitask learning component, we engineered custom task-specific regression heads for the ESM-2 model (**Fig. S1A**). On the embedding layer of ESM-2, a linear layer with dimensions equal to the number of tasks was set as task-specific heads. Additionally, an intermediate linear layer with 252 dimensions connecting the embedding layer and the task-head layer was set.

In CoVFit, a single input sequence is linked to multiple target variables due to the multitask learning framework. For example, regarding DMS data, a typical S protein mutant (input sequence) is linked to relative binding affinity values for >1,000 mAbs (target variables). To boost computational efficiency, CoVFit utilizes an architecture that processes a single input sequence alongside its multiple corresponding target variables in parallel, rather than processing pairs of the same input sequence and one target variable sequentially (**Fig. S1B**). As a result, the loss values for multiple target variables associated with a single input sequence are calculated simultaneously.

However, the number of tasks linked to each input sequence can differ greatly, especially when comparing the variant S protein sequences used for fitness prediction (up to 17 tasks) against the mutant sequences used for DMS predictions (up to 1,548 tasks). Consequently, the magnitude of the loss value for each dataset can vary significantly based on the number of associated tasks, which can lead to training instability. To stabilize the training process, CoVFit utilized non-overlapping random sampling to create data chunks where a single input sequence is associated with target variables for a maximum of 10 tasks. These generated sequence-variable chunks were then used as the training inputs.

For the loss function, CoVFit utilizes a custom least squares approach weighted according to individual tasks. In principle, the weights were determined to be proportional to the reciprocals of the task frequencies. One exception was implemented where for fitness prediction tasks for genotypes emerged after January 1, 2022, we adjusted the weights by doubling them.

### Model finetuning and performance evaluation

In CoVFit, we finetuned the custom model based on ESM-2 using the LoRA technique implemented in the Hugging Face PEFT v.0.5.0. Low rank adaptors were injected into the weight matrices of the key, query, and value components, as well as those for the dense layers. Full finetuning was applied to these adaptors and the custom regression heads added onto ESM-2, while the other, original layers were kept frozen in their pretrained state. A rank parameter of r=8 and a scale parameter of alpha=16 were used. Consequently, out of the total 659,741,475 parameters, the model has 7,768,974 trainable parameters, which constitutes approximately 1.18% of the total. The LoRA dropout rate was set at 0.05.

For finetuning, the AdamW optimizer was used with a weight decay parameter of 0.02. The maximum learning rate was set at 2.0E-4 with a linear learning rate scheduler, and the training was conducted over 30 epochs. The batch size was set at 4 with gradient accumulation steps of 2.

The genotype–fitness and DMS datasets were randomly divided into training, evaluation, and test datasets in a 6:2:2 ratio. For the genotype–fitness dataset, we considered the combinations of country and Nextclade clade, ensuring that data representing each combination were evenly distributed across the training-evaluation and test datasets. Similarly, for the DMS dataset, the types of mAbs were considered during the data splitting process. We conducted the data splitting with a five-fold cross-validation approach.

In our experiments aimed at assessing the model’s ability to predict the performance of future variants, we began by dividing the genotype–fitness dataset into two: one for past variants and another for future variants, based on their emergence dates. For experiments focusing on the BQ.1, XBB, CH.1.1, and BA.2.86 lineages, we set the cutoff date as September 1, 2022. We then excluded these specific target lineages from the past variant dataset. Similarly, for the experiment that specifically targets the BA.2.86 lineage, we chose August 1, 2023, as the cutoff date and removed this lineage from the past dataset. After this step, we further divided the past dataset into three subsets for training, validation, and testing purposes, following the methodology described in the paragraph above.

In our experiments designed to assess the importance of including DMS data for immune evasion, we trained alternative instances without incorporating the DMS dataset for comparison. Likewise, in our experiments evaluating the significance of the domain adaptation step, we employed the original ESM-2 model rather than the version adapted to the coronaviral S protein dataset. In these experiments, the past–future dataset with a cutoff date of August 31, 2022, was used.

### CoVFit-CLI

The CoVFit-CLI tool packages CoVFit_Nov23_ via pyinstaller 6.4.0 using Python v.3.10.9, torch v.2.1.2, transformers 4.37.1, and bio v.1.5.9 with CUDA v.12.3 on x86_64 Linux, kernel 5.15.0.

### Development of fitness prediction models based on non-deep learning models

We constructed fitness prediction models based on LASSO, Random Forest, and LightGBM to compare the prediction performance of CoVFit with those of these models. LASSO employs a linear regression framework enhanced with L1 regularization, offering a method to include penalty terms that reduce overfitting by shrinking some coefficients to zero. In contrast, Random Forest and LightGBM are advanced, decision tree-based models known for their greater expressive capability. These models aim to predict a variant’s fitness value based on its amino acid mutation profile in the S protein and the country of origin. Both the mutation profile and the country data were one-hot encoded to serve as input features for the models.

We trained these models and evaluated their performance using the past–future variant datasets with a cutoff date of August 31, 2022, and the five-fold cross validation scheme as described the **Model Finetuning and performance evaluation** section. In the training dataset, we selected 200 features to be used as inputs of the models according to the feature importance estimated by Random Forest. We trained the models with hyperparameter-tuning using a Bayesian optimization method. In this process, R^2^ was used as the optimization metric, and the number of iterations was set at 20. The parameter spaces searched in this step are described in detail in the GitHub repository (https://github.com/TheSatoLab/CoVFit).

The machine learning models above were reconstructed using Python v.3.9.13, pandas v.1.4.4, numpy v.1.21.5, lightgbm v.3.3.5, scikit-learn v.1.0.2, and scikit-optimize v.0.9.0.

### Phylogenetic analysis

We created the dataset for phylogenetic analysis as a subset of the dataset of the representative viral genome sequences encoding respective S protein genotypes (EPI_SET_ID: EPI_SET_240311ma; **see Preparation of genotype– fitness dataset**). We removed sequences matching the following criteria: i) sequences with >3% ambiguous characters across positions 265 to 29,673 (in alignment with the Wuhan-Hu-1 reference (GenBank accession number: NC_045512.2)) and ii) sequences classified as “recombinant” according to Nextclade clade assignments. Additionally, we included the Wuhan-Hu-1 reference genome sequence to our dataset. The dataset for viral genome sequences used in the phylogenetic analysis, except for the Wuhan-Hu-1 reference genome, are summarized under the EPI_SET_ID: EPI_SET_240311rk, which can be accessed through the GISAID website (https://gisaid.org/).

The nucleotide viral genome sequences were aligned to the reference sequence of Wuhan-Hu-1 using Minimap2 v.2.17^45^. This alignment was then converted into a multiple sequence alignment following the GISAID phylogenetic analysis pipeline (https://github.com/roblanf/sarscov2phylo). Sites corresponding to positions 1–265 and 29,674–29,903 in the reference genome were masked, that is, converted to ‘NNN’, to exclude them from subsequent analyses. The maximum likelihood phylogenetic tree was constructed using IQ-TREE v.2.1.4_beta, adopting the GTR+I+G nucleotide substitution model^46^. To assess the reliability of the phylogenetic tree nodes, an ultrafast bootstrap analysis was performed with 1,000 replicates. A time-resolved phylogenetic tree was inferred from the constructed tree using TreeTime v.0.11.1, with the rerooting strategy set to ‘oldest’^47^, resulting rerooting by the Wuhan-Hu-1 strain. The S protein sequences for ancestral nodes were reconstructed also using TimeTree with the default options.

### Detection of phylogenetic branches with fitness elevation using CoVFit

To infer the impact of mutations on fitness through the observed evolution of SARS-CoV-2, we analyzed the increase in predicted fitness across all branches of the SARS-CoV-2 phylogenetic tree, as outlined in the **Phylogenetic analysis** section. We employed five CoVFit_Nov23_ models, developed via a five-fold cross-validation, to predict the fitness values for both existing and reconstructed ancestral S protein sequences within the tree. Since CoVFit predicts fitness across multiple countries, we averaged these predictions to obtain a single representative fitness value for each sequence, resulting in five representative fitness values per sequence. We compared these values between each node and its ancestral node, calculating the mean fitness gain for the branches connecting them. Statistical significance of fitness changes was determined using Welch’s t-test, with multiple testing correction applied via the Benjamini-Hochberg method. Branches with a false discovery rate (FDR) less than 0.1 were considered statistically significant. We also identified mutations in the S protein acquired along each branch by comparing the S protein sequences at both ends of the branch. The detected fitness elevation events are summarized in **Table S3**.

### Characterization of the F456L substitution using publicly available DMS data

Position-wise scores for escape from humoral immunity were calculated using escape estimator^32^ based on DMS data for the ability to evade mAbs presented in Cao et al.^29^ (https://github.com/jbloomlab/SARS2_RBD_Ab_escape_maps) (shown in **Fig. 5C**). For the ACE2 binding and protein expression DMS data, we retrieved the per-site variant score results presented by Taylor and Starr^31^ (https://github.com/tstarrlab/SARS-CoV-2-RBD_DMS_Omicron-XBB-BQ/blob/main/results/final_variant_scores/final_variant_scores.csv). We filtered for mutant L on position 456 and retrieved the ‘delta_bind’ and ‘delta_expr’ values (representing the mean of values across replicates minus the mean for the reference residue) for each variant target (shown in **Fig. 5D**).

### CoVFit-based *in silico* (deep) mutational scanning analysis

Instances of the CoVFit_Nov23_ model were utilized to infer the fitness of S protein mutants. Frist, the mean fitness value for each mutant was calculated across different countries. Subsequently, these mean values were averaged across all five CoVFit_Nov23_ model instances, yielding a singular average fitness value for each S protein mutant. This streamlined fitness value was then compared to the fitness of the original backbone S protein sequence.

### Epidemic analysis on JN.1 subvariants

We retrieved all SARS-CoV-2 genome sequences and their associated metadata available up to March 11, 2024, from GISAID (https://gisaid.org/). To ensure data quality, sequences were excluded from analysis based on the following criteria: (i) absence of collection date; (ii) samples taken from animals other than humans; (iii) more than 2% undetermined nucleotides; or (iv) samples collected during quarantine. Our focus was on the JN.1 lineages, specifically those tagged as “VOI GRA (JN.1+JN.1.1*)”, collected between December 1, 2023, and February 29, 2024 (EPI SET ID: EPI_SET_240315fc). We categorized the JN.1 lineages into five groups: JN.1 with F456*, JN.1 with R346*, JN.1 with K478*, JN.1 with both F456* and R346*, and other JN.1 lineages, where an asterisk denotes an arbitral amino acid type. The frequencies of these five JN.1 categories were calculated in 7-day intervals for each geographic region within the JN.1 population. Data for Africa was excluded due to a low total sequence count (n = 216).

### Methodological Discussion: Room for improvement of CoVFit

We recognize the presence of multiple areas where CoVFit could potentially be improved with future development. First, since our current model is solely trained on S protein sequences, it may be possible to improve its performance by including information on additional viral proteins. Previous studies have identified mutations associated with increased fitness also in non-S proteins, particularly in the nucleocapsid (N) protein, supporting the possible effectiveness of this approach^6^. In the current setting, the effects of mutations in non-S proteins are absorbed into the effects of mutations in the S protein that have linkage disequilibrium relationships with these mutations. However, it is certain that the S protein has a particularly strong impact on fitness compared to other viral proteins^6^. Therefore, it is unclear to what extent prediction accuracy would be improved by adding information from other viral proteins. There is even a possibility that generalizability could be decreased by including other viral proteins due to the decrease of signal-to-noise ratio. Given that mutations enhancing viral fitness are predominantly found in the S1 subunit of the S protein, focusing solely on the S1 subunit as an input may result in a higher signal-to-noise-ratio, providing a more effective approach.

Second, it may also be possible to improve the performance by including various DMS data, such as those on other viral phenotypes. In this study, we only used DMS data on the immune evasion ability against mAbs. However, given the significant impact of the ACE2 binding affinity of the S protein on fitness, employing DMS data for this trait could improve predictive performance. In our preliminary experiments, however, we found that the convergence speed for DMS data on binding affinity to ACE2 was much slower compared to other tasks. Considering the difficulty of simultaneous learning, we decided not to use this DMS data in this study. Similarly, while we used DMS data based on the ancestral strain (D614G), incorporating DMS data obtained by experiments using S proteins of other variants (particularly variants emerged recently) could potentially further improve predictive performance.

The third consideration is the scaling up of the model. It is generally known that language models improve in performance as they increase in size. Indeed, benchmark tests using ESM-2 have shown that changing the model size from the 650M, used in this study, to 3B or 15B can lead to slight improvements in performance regarding the prediction of protein structures^27^. For our method, using larger models like 3B or 15B models may also enhance performance. Recently developed techniques, such as QLoRA, a quantization-based LoRA^48^, make it possible to finetune even 15B models in a limited GPU resource. However, we faced issues with incompatibility between CoVFit and QLoRA, leading us to abandon the development of a QLoRA-based model. It is also important to note that scaling up the model can significantly increase training and inference times.

The fourth consideration is data augmentation for fitness data. In viral genome surveillance, there can be a delay of several weeks to months between the date of sample collection and the date of data submission to databases. Furthermore, in viral genome surveillance, the number of viral genomes newly sequenced have gradually decreased in recent years (https://gisaid.org/). Consequently, the most recent genomic data tends to be under-sampled. However, this recent data is considered to contain more critical information for predicting future variants compared to older data. Therefore, increasing the proportion of recent data in the training dataset or employing data augmentation techniques, which artificially expand the dataset, might enhance the model’s ability to generalize to future variants.

Lastly, since we have not conducted an exhaustive investigation of the model’s hyperparameters, the model’s performance could be improved by adjusting them. Adjustable hyperparameters include data normalization methods, network architecture, task weight balance, the optimizer algorithm, learning rate, and maximum epoch numbers.

## Author Contributions

Jumpei Ito designed the study, the main conceptual ideas, and the proof outline. Adam Strange developed ESM-2_Coronaviridae_ and the CovFit-CLI standalone command line tool.

Wei Liu constructed a series of non-deep learning prediction methods. Gustav Joas collected and created the *Coronaviridae* dataset.

Spyros Lytras performed detailed analysis on the F456L substitution.

Jumpei Ito performed the other parts of CoVFit development and computational analyses.

Jumpei Ito made the figures and wrote the initial draft of the manuscript. Adam Strange and Spyros Lytras provided editing.

Kei Sato and The Genotype to Phenotype Japan (G2P-Japan) Consortium contributed to the project administration.

All authors reviewed and proofread the manuscript.

## Acknowledgments

We gratefully acknowledge all data contributors, i.e. the Authors and their Originating laboratories responsible for obtaining the specimens, and their Submitting laboratories for generating the genetic sequence and metadata and sharing via the GISAID Initiative, on which this research is based. We would like to thank all members belonging to The Genotype to Phenotype Japan (G2P-Japan) Consortium. The super-computing resources were provided by Human Genome Center and the Information Technology Center, The University of Tokyo.

This study was supported in part by JST PRESTO (JPMJPR22R1, to Jumpei Ito); JSPS KAKENHI Grant-in-Aid for Early-Career Scientists (23K14526, to Jumpei Ito); AMED SCARDA Japan Initiative for World-leading Vaccine Research and Development Center “UTOPIA” (JP223fa627001, to Jumpei Ito; JP223fa627001, to Kei Sato); AMED ASPIRE (JP23jf0126002, to Kei Sato); AMED SCARDA Program on R&D of new generation vaccine including new modality application (JP223fa727002, to Kei Sato); AMED Research Program on Emerging and Re-emerging Infectious Diseases (JP22fk0108146, to Kei Sato; JP21fk0108494 to G2P-Japan Consortium, and Kei Sato; JP21fk0108425, to Kei Sato; JP21fk0108432, to Kei Sato; JP22fk0108534, to Kei Sato; JP22fk0108511, to Kei Sato); AMED Research Program on HIV/AIDS (JP22fk0410039, to Kei Sato); JST CREST (JPMJCR20H4, to Kei Sato); JSPS KAKENHI Fund for the Promotion of Joint International Research (International Leading Research) (JP23K20041 to Kei Sato); JSPS Core-to-Core Program (A. Advanced Research Networks) (JPJSCCA20190008, to Kei Sato); The Tokyo Biochemical Research Foundation (to Kei Sato); and Mitsubishi Foundation (to Kei Sato); JSPS International Research Fellow (to Gustav Joas).

## Consortia

### The Genotype to Phenotype Japan (G2P-Japan) Consortium

Keita Matsuno^11^, Naganori Nao^11^, Hirofumi Sawa^11^, Keita Mizuma^11^, Isshu Kojima^11^, Jingshu Li^11^, Tomoya Tsubo^11^, Shinya Tanaka^11^, Masumi Tsuda^11^, Lei Wang^11^, Yoshikata Oda^11^, Zannatul Ferdous^11^, Kenji Shishido^11^, Takasuke Fukuhara^11^, Tomokazu Tamura^11^, Rigel Suzuki^11^, Saori Suzuki^11^, Shuhei Tsujino^11^, Hayato Ito^11^, Yu Kaku^1^, Naoko Misawa^1^, Arnon Plianchaisuk^1^, Ziyi Guo^1^, Alfredo A Hinay Jr.^1^, Kaoru Usui^1^, Wilaiporn Saikruang^1^, Keiya Uriu^1^, Yusuke Kosugi^1^, Shigeru Fujita^1^, Jarel Elgin M.Tolentino^1^, Luo Chen^1^, Lin Pan^1^, Wenye Li^1^, Mai Suganami^1^, Mika Chiba^1^, Ryo Yoshimura^1^, Kyoko Yasuda^1^, Keiko Iida^1^, Naomi Ohsumi^1^, Shiho Tanaka^1^, Kaho Okumura^1^, Kazuhisa Yoshimura^12^, Kenji Sadamas^12^, Mami Nagashima^12^, Hiroyuki Asakura^12^, Isao Yoshida^12^, So Nakagawa^13^, Akifumi Takaori-Kondo^14^, Kotaro Shirakawa^14^, Kayoko Nagata^14^, Ryosuke Nomura^14^, Yoshihito Horisawa^14^, Yusuke Tashiro^14^, Yugo Kawai^14^, Kazuo Takayama^14^, Rina Hashimoto^14^, Sayaka Deguchi^14^, Yukio Watanabe^14^, Yoshitaka Nakata^14^, Hiroki Futatsusako^15^, Ayaka Sakamoto^14^, Naoko Yasuhara^14^, Takao Hashiguchi^14^, Tateki Suzuki^14^, Kanako Kimura^14^, Jiei Sasaki^14^, Yukari Nakajima^14^, Hisano Yajima^14^, Takashi Irie^15^, Ryoko Kawabata^15^, Kaori Sasaki-Tabata^16^, Terumasa Ikeda^17^, Hesham Nasse^17^, Ryo Shimizu^17^, MST Monira Begum^17^, Michael Jonathan^17^, Yuka Mugita^17^, Sharee Leong^17^, Otowa Takahashi^17^, Kimiko Ichihara^17^, Takamasa Ueno^17^, Chihiro Motozono^17^, Mako Toyoda^17^, Akatsuki Saito^18^, Maya Shofa^18^, Yuki Shibatani^18^, Tomoko Nishiuchi^18^, Jiri Zahradni^19^, Prokopios Andrikopoulos^19^, Miguel Padilla-Blanco^19^, Aditi Konar^19^

^11^ Hokkaido University, Sapporo, Japan

^12^ Tokyo Metropolitan Institute of Public Health, Tokyo, Japan

^13^ Tokai University, Kanagawa, Japan

^14^ Kyoto University, Kyoto, Japan

^15^ Hiroshima University, Hiroshima, Japan

^16^ Kyushu University, Fukuoka, Japan

^17^ Kumamoto University, Kumamoto, Japan

^18^ University of Miyazaki, Miyazaki, Japan

^19^ Charles University, Vestec-Prague, Czechia

## Data availability

Surveillance datasets of SARS-CoV-2 genomes are available from the GISAID database (https://www.gisaid.org; EPI_SET_240307pq; EPI_SET_240311ma; EPI_SET_240311rk; EPI_SET_240315fc). The supplemental table for each GISAID dataset is available in the GitHub repository (https://github.com/TheSatoLab/CoVFit).

## Code availability

The computational codes used in this study are available in the GitHub repository (https://github.com/TheSatoLab/CoVFit).

## Figure Legends

**Fig. S1 CoVFit architecture.**

A) Architecture of CoVFit. The input of CoVFit is the amino acid sequence(s) of SARS-CoV-2 S protein, aligned to the reference Wuhan-Hu-1 strain’s S protein. The input is tokenized by the ESM-2 tokenizer with truncation to the first 1–1,024 amino acids. The output of CoVFit is predicted outcomes for respective tasks, including relative fitness value for each country and escape score for each mAb. For multitask learning, the task-specific regression dense layer is set on the embedding out layer of ESM-2 with an intermediate dense layer in-between. ESM-2 layers are finetuned with LoRA, an efficient finetuning technique^42^, while the intermediate and task-specific regression layers are fully finetuned.

B) Training scheme for efficient multitask learning in CoVFit. CoVFit processes a single input sequence alongside its multiple corresponding target variables (up to 10 tasks, selected by non-overlapping random sampling) in parallel, rather than processing pairs of the same input sequence and one target variable sequentially. Consequently, the loss values for multiple target variables associated with a single input sequence are calculated simultaneously. This scheme can reduce the redundant calculations of self-attention for a given input sequence, leading to saving computational time.

C) Perplexity scores of ESM-2 and ESM-2_Coronaviridae_ models, which represent the prediction performance for masked learning tasks. Lower score indicates better performance. The score was calculated for two protein sequence datasets: i) the SARS-CoV-2 S protein (future variants) dataset, including SARS-CoV-2 S protein genotypes emerged later than August 31, 2022, which are not included in the training data for domain adaptation. ii) A subset of UniRef50, representing a variety of protein sequences including proteins other than the S protein. See **Domain adaptation** in **Methods** for detail.

**Fig. S2 Datasets used for CoVFit training.**

A) Estimated relative fitness (R_e_) of each S protein genotype in each country. X-axis indicates the emergence date of each genotype. Dot color indicates the Nextclade clade classification.

B) UMAP plot representing the similarity of DMS results among mAbs. Dot represents a specific mAb. Dot color indicates the epitope class of each mAb.

C) Mean escape score over a mutation across mAbs. Results are stratified according to the epitope class.

D) Immune evasion capacity of major variants. Log_10_(-IC_50_) values for 1,548 mAbs against variants are shown. A lower log_10_(-IC_50_) signifies reduced mAb binding (indicating enhanced escape capability of a variant). Top) Log_10_(-IC_50_) value distribution with the mean value (dot). Middle) Log_10_(-IC_50_) values for individual mAbs. Bottom) Mean log_10_(-IC_50_) value per epitope class. Data sourced from Cao et al.^29^.

**Fig. S3 Prediction performance of CoVFit.**

A) Scatter plot for mAb escape prediction, aggregating results from five-fold cross-validation. Results are categorized by epitope group. Dot density is represented by color. The escape score was scaled so that 0 and 95 percentiles fell within the range 0–1, and values above 95 percentile were clipped to 1. A dashed line with a slope of 1 and intercept of 0 is shown.

B) Spearman’s correlation scores for fitness prediction in each country.

C) Spearman’s correlation scores for predicting mAb neutralization escape scores in each type of mAb.

**Fig. S4 Prediction performance of CoVFit for BA.2.86.**

A) Strategy to evaluate prediction performance for future variants including BA.2.86 and its descendant, JN.1. Model instances, referred to as CoVFit_Sep23_, were trained using data for variants emerged by September 30, 2023.

B) Comparison of predicted fitness among major variants. Predicted fitness value in each country (**top**), mean observed fitness value across countries (**middle**), and number of variant sequences included in the past dataset (**bottom**), are shown.

**Fig. S5 Comparison of prediction performance among methods.**

A) Scatter plot for fitness prediction, aggregating results from five-fold cross-validation. Both past (light gray) and future (gray) variants are included. Regarding future variants, the mean prediction across five-fold cross-validation datasets is shown. A dashed line with a slope 1 and intercept 0 is shown.

B) Scatter plot for fitness prediction including only future variants. Mean (dot) and standard deviation (error bar) across prediction results by five-fold model instances are shown. Dot color denotes the Nextclade clade classification. In addition to the line with slope 1 and intercept 0 (black), the estimated linear regression line (gray) is shown.

**Fig. S6 Detection of fitness elevation events during the evolution of SARS-CoV-2, including pre-Omicron and Omicron lineages.**

A) Phylogenetic tree of SARS-CoV-2 variants. Trees shown in Figs. 4 and 5 are the subtree of this tree.

B) Inference of change in fitness through SARS-CoV-2’s evolution. Branch color indicates an inferred fitness value for each phylogenetic node, including both observed and reconstructed ancestral genotypes of S proteins in the phylogenetic tree.

C) Detection of fitness elevation events during SARS-CoV-2’s evolution. Dot color indicates inferred fitness gain, calculated as difference in predicted fitness between a node and its parental node.

D) Mean fitness gain over a specific mutation during SARS-CoV-2 evolution. Since some mutations have been acquired multiple times, the mean value of fitness gain among acquisition events was used as the “fitness gain [per mutation]” score.

E) Enrichment of fitness-associated mutations in the RBD, particularly in its RBM.

F) Mapping the site-wise fitness gain value on the 3D structure of the ancestral D614G S protein (PDB: 7BNN)^49^. If multiple mutation types are present in a specific site, the maximum value is shown as the “fitness gain [per site]” score. The plot was generated using Chimera X^50^.

## Supplemental Tables

**Table S1 Estimated relative R_e_ of each S protein genotype in each country. Table S2 List of sequences used in domain adaptation process.**

**Table S3 Identification of phylogenetic branches where significant fitness elevation events were detected.**

